# Aberrant placental structure is corrected with repeated nanoparticle-mediated IGF1 treatments in a guinea pig model of fetal growth restriction

**DOI:** 10.1101/2025.06.05.658039

**Authors:** Baylea N. Davenport, Rebecca L. Wilson, Alyssa A. Williams, Jaimi A. Gray, Edward L. Stanley, Helen N. Jones

## Abstract

Fetal growth restriction (FGR) is most commonly due to placental insufficiency. There are currently no treatments for placental insufficiency or FGR, and the only intervention is iatrogenic pre-term delivery. We have previously shown efficacy of repeated placental nanoparticle-mediated *insulin-like 1 growth factor* (*IGF1*) treatment in improving placental efficiency (increased fetal-placental weight ratio) and correcting fetal growth in a maternal nutrient restriction (MNR) guinea pig model of FGR. Here we investigate the structural remodeling of the placenta following 3 repeated intraplacental injections of nanoparticle-mediated *hIGF1* that may underpin the published improvements in placental efficiency and fetal growth. We hypothesize placenta structural changes (reduced exchange area, altered vascular structure) that we and others have previously shown in the FGR/MNR placenta which lead to deficits in placental function are mitigated by our repeated nanoparticle-mediated *hIGF1* treatment. Sham-treated MNR placentas displayed disorganized microvasculature labyrinthine exchange areas with a reduction in placental capillary number and maternal blood space areas, and an increase in the volume of the placenta macrovasculature. Repeated nanoparticle-mediated *hIGF1* treatment, however, resulted in an improved exchange area with normalized placental capillary number and maternal blood space area, and macrovasculature volume. This data demonstrates repeated nanoparticle-mediated *hIGF1* delivery corrects aberrant placenta structure likely leading to improved gas exchange and transfer of nutrients to the fetus restoring fetal growth.

## INTRODUCTION

Fetal growth restriction (FGR) affects 10% of humans worldwide and is defined as a fetus failing to reach their growth potential [1-3]. While diagnosis criteria varies between regions, the most commonly accepted criteria according to ACOG (American College of Obstetricians and Gynecologists) guidelines is an estimated fetal weight (EFW) or abdominal circumference below the 10th percentile for gestational age. FGR can lead to stillbirth and iatrogenic preterm delivery, with FGR neonates having increased risk of developing numerous comorbidities linked to aberrant fetal programming [4]. FGR is associated with lifelong consequences stemming from the developmental origins of health and disease (DOHaD) including cognitive delays/ deficits, cardiovascular disorders, and metabolic disorders [5, 6]. Most often, the cause of this condition is due to placental insufficiency: the placenta’s failure to supply sufficient oxygen and nutrients to the fetus for proper growth and development [7, 8].

Placental functions include gas/oxygen exchange, immune barrier, endocrine function, nutrient transfer, and waste excretion and is vital for fetal growth and development. Proper blood flow into, and structure of, the placenta is critical to deliver all necessary oxygen and nutrients from the maternal circulation, across the syncytium and villous core to the fetal circulation. The macrovasculature structures, or the stem-like structures branching from either the uteroplacental vessels (of the maternal circulation) and the umbilical cord (the fetal circulation) lead into highly branched microvascular exchange areas to support the increasing transfer and exchange throughout pregnancy. Insufficient placentas of growth restricted fetuses display abnormalities in both the utero-placental circulation and the feto-placental circulation, impeding these pivotal processes [9, 10]. Human FGR studies show maternal vascular abnormalities in spiral artery remodeling within the uterus leading to increased resistance shown by uterine artery doppler [9-13]. The fetoplacental vascular network also displays increased resistance with underdeveloped terminal villi/hypovascularity [14-16]. These aberrant vasculatures restrict blood flow and reduce gas exchange, waste exchange, and nutrient transport in the microvasculature exchange area [9, 10, 14].

Guinea pigs offer one of the most useful models to study placental development and fetal growth restriction due to their developmental similarities to humans [17]. Like humans, guinea pigs have invasive, haemomonochorial, discoid placentas. Similar to the human placenta which is organized into cotyledons each being supplied by a single spiral artery from the maternal circulation, the guinea pig placenta has distinct lobules of circulatory exchange with microvasculature bringing maternal blood to the syncytial lined maternal lacunae of each lobules exchange region.[12]. Similar to the chorionic plate vessels and stem villi vessels in the human, the feto-placental circulation macrovasculature of each lobule branches to form a network of microvasculature. Bloodflow within the maternal lacunae and apposing macrovasculature carrying fetal blood is counter-current to maximize the area of nutrients and oxygen exchange[18]. Guinea pig fetuses recapitulate human developmental milestones throughout gestation and after birth making them an excellent candidate to study both placental and fetal development [17, 19]. To model FGR we employed the well-established maternal nutrient restriction (MNR) model [20]. The MNR model uses a reduction in food intake (70% of Control) for the mother to increased stress/cortisol [19-21]. Roberts et al. previously showed at both mid and late gestation, the MNR guinea pig placenta had reduced exchange surface area and increased barrier thickness, recapitulating the dysregulated microvasculature exchange area seen in human placentas [22]. This model has been consistently shown to induce insufficient placental development and function, and impact similar signaling cascades that lead to reduced growth in human cases of FGR [20-22].

We have employed the guinea pig MNR model to study the effects of our novel gene therapy for the correction of placental insufficiency and in turn FGR [23, 24]. The insulin-like growth factor (IGF) axis is one of the key signaling cascades shown to be downregulated in placentas of FGR humans and animal models [25, 26]. IGF1 is produced by placental trophoblast throughout gestation with numerous roles in regulating nutrient transport, proliferation, and angiogenesis [27, 28]. Using ultrasound guided intraplacental injection of nanoparticles containing the *hIGF1* gene under the control of a trophoblast-specific primer, we previously demonstrated [29-31] increased placental capillary volume density and reduced interhaemal distance between maternal and fetal blood supply at mid-pregnancy, which may facilitate increased gas exchange and nutrient transfer [30]. With three injections starting after establishment of FGR (each 8 days apart) fetal growth was corrected to Control weight by term. [23].

The overall aim of this study was to investigate the structural changes of the placenta labyrinth in a model of placental insufficiency and assess the impact of repeated nanoparticle-mediated trophoblast-specific *hIGF1* treatments. We hypothesize that our treatment improved placenta efficiency by restoring placental labyrinth vasculature structure (both micro and macro) for improved blood flood, gas exchange, and access of syncytium for nutrient uptake.

DiceCT has been a critical tool enabling comparative biologists to increase their knowledge on taxonomy and museum archivists to create digital databases of museum specimens. The oVert (OpenVertebrate) project was designed to provide open access digital 3D vertebrate anatomy models to researchers and the public using CT for skeletal models and diceCT imaging for soft tissue models [32, 33]. We partnered with the oVert team at the Florida Museum of Natural History to use this technology for our pre-clinical work on FGR. Using this technique, we sought to 3D render the macrovasculature of the guinea pig placenta to understand the vascular changes within the placental lobules during placental insufficiency and following nanoparticle-mediated IGF1 treatment. From this segmentation we aimed to evaluate the size (volume/surface area) of the lobule vasculature networks to understand the potential impact on blood flow/gas exchange in MNR and using immunohistochemistry to evaluate the labyrinth maternal lacunae and microvascular changes following treatment.

## METHODS

### Nanoparticle Synthesis

Nanoparticles were complexed using a lyophilized non-viral PHPMA_115_-b-PDMEAMA_115_ co-polymer reconstituted in water combined with plasmid containing the *hIGF1* gene under the control of a trophoblast-specific promoter, *CYP19A1*. 50 μg of plasmid was suspended in a 200 μL volume solution at room temperature. Detailed methods of copolymer synthesis and nanoparticle formation can be found in Wilson et al., 2022 [30].

### Animal Husbandry and Mating

Animal care and usage was approved by the Institutional Animal Care and Usage Committee at the University of Florida (Protocol #202011236). Female Dunkin-Hartley guinea pigs (Dams) were purchased from Charles River Laboratories (Wilmington, MA) at 500–550 g (∼ 8-9 weeks of age). Animals were housed in an environmentally controlled room (22°C/72°F, 50% humidity 12 h light-dark cycle). Upon arrival, food (LabDiet diet 5025: 27% protein, 13.5% fat, and 60% carbohydrate as % of energy) and water were provided ad libitum. Guinea pigs were acclimatized for 2 weeks prior to being assigned to either ad libitum diet (termed Control: n = 6) or maternal nutrient restriction (MNR) diet (n=12). Assignment was done by ranking animals heaviest to lightest and systematically assigning them to each group for even biological weight distribution. MNR diet consists of a 70% food intake diet based on kilogram of body weight of control from 4 weeks pre-mating through mid-pregnancy (GD35), then increased to 90% food intake through term to maintain pregnancy as previously published [30].

### Intraplacental Nanoparticle Injections, Animal Sacrifice, and Tissue Collection

Ultrasound guided intraplacental nanoparticle delivery was performed a total of 3 times, each 8 days apart from mid pregnancy till term (GD36,44,52±3) with guinea pigs being sacrificed 8 days after the final injection (GD60±3). One placenta per litter was injected with either nanoparticles containing Cyp19A-*hIGF1* (MNR + IGF1 n=6) or a non-expressing plasmid sham nanoparticle (Control n=6, MNR n=6). In the MNR + IGF1 dams, placentas were separated based on receiving a direct injection of nanoparticle-mediated *hIGF1* “MNR + IGF1 (Direct Injection)” or by being indirectly exposed to circulating residual nanoparticle-mediated *hIGF1* “MNR + IGF1 (Indirect Exposure)” which was further confirmed by expression levels of *hIGF1* in each placenta via qPCR [23]. Dams were sacrificed via carbon dioxide asphyxiation followed by cardiac puncture and exsanguination. Fetuses (Control: n=8 female and n=11 male, MNR: n=5 female and 11 male, MNR + IGF1: n=6 female and 10 male) and the delivered placenta (placenta, sub-placenta, and decidua) were removed from the uterus and weighed. Fetal sex was determined at this time by examination of the gonads. Here it was determined that by random chance only male fetuses received direct injections (as we cannot reliably determine fetal sex at time of first injection). Blood was collected via cardiac puncture from dams and fetuses at time of sacrifice. To reduce bias, all subsequent analyses were performed blinded. For in-depth synopsis of injections, litter size, etc see Davenport, Wilson, et al 2024 [23].

### Placenta Preparation

At time of sacrifice all placentas were halved (each half included placenta, sub-placenta, and decidua still attached). One half was fixed in 4% Paraformaldehyde for 48 hours and washed in PBS for 24 hours before being placed in increasing ethanol concentrations until reaching 70% ethanol. Placenta specimen remained in 70% ethanol until staining for diceCT (diffusible iodine contrast enhanced computed tomography) or being process for histological analysis [34].

### Iodine Staining

In placentas analyzed for DiceCT, iodine staining was performed by combining 15% Lugol’s iodine (Sigma) with 2x Sorenson’s buffer [32, 35]. Placentas were completely submerged in 1.25% Lugol’s solutions for 7 days, followed by 2.5% Lugol’s solution for 10 days. DiceCT introduces X-ray opaque iodine into soft tissues via diffusion, and iodine in the form of triiodide binds to lipids and carbohydrates abundant in soft tissue structures. The staining process was considered complete when the iodine in the container remained a deep red color. Any tissues considered overstained were placed briefly into a water bath to allow the iodine to partially diffuse back out of the tissue. N numbers: Control: Dams: n=2, placentas n=6, MNR: Dams: n=3 female, placentas n=9, MNR + IGF1: Dams: n=3, placentas n=9

### DiceCT Scanning

Placentas were removed from iodine, individually wrapped in packing foam, and placed in small plastic bags. Plastic bags were rolled and placed in small plastic tubes to reduce movement throughout the scans. Space was left empty in the bottom of the plastic bag below the specimen to provide room for any residual iodine to drain for optimal scanning without scattering or saturation of the light source from pooled iodine. Aluminum wire was placed in each tube beside each specimen to serve as a material of known density for gray scale standardization at a later step in the analysis. The GE Phoenix c| tome | x m was used to perform nano-CT imaging. Scans were performed at 80 kV and 200 uA, with 1499 second exposure timing, 3x frame averaging with 1 skipped frame, a gain (sensitivity) value of S=2, and a 0.3mm aluminum filter with a voxel size of ∼0.016 in the x, y, and z direction. Raw tomography projections were reconstructed using GE Datos R software (General Electric, version 2.8.2) to generated tiff (tagged image file format) stacks of approximately 2000 cross-sectional images. 3D reconstructions of placentas imported into Volume Graphics (VG) Studio Max version 2024.2.1 (Volume Graphics, Heidelberg, Germany) for segmentation and analysis.

### DiceCT Analysis/Segmentation

3D reconstructions were imported into VG Studio Max and aluminum wires were used to standardize gray scale values. The “paint and segment” machine learning tool was used to identify packaging/air surrounding the tissue, placental tissue, and the macrovasculature within the placenta. To avoid overtraining the machine learning models, training was performed for no longer than 30 minutes. Once segmentation was rendered from the machine learning, the “draw” function was used to manually clean-up any identification errors from knowledge of placenta morphology.

### Immunohistochemistry

Remaining fixed placentas not used for DiceCT were processed and paraffin embedded as standard. 5 μm thick sections were cut and slide-mounted for double-label immunohistochemistry (IHC). Sections were de-waxed and rehydrated using histo-clear and ethanol following standard protocols. Antigen retrieval was performed with 0.03% protease (Sigma) at 37°C for 15 minutes. Endogenous peroxidase activity was blocked with 3% hydrogen peroxide for 30 minutes before blocking with serum-free protein block (Dako) for 10 minutes. Placental capillary endothelium was identified using anti-vimentin (Dako Vim3B4, 1:100 [0.5 μg/ml]) and placental trophoblasts were identified using anti-pan cytokeratin (Sigma C2562, 1:200 [10 μg/ml]). Anti-vimentin antibody was diluted in 10% guinea pig serum with 1% BSA and applied to slides overnight at 4°C. Sections were washed in PBS before biotinylated anti-mouse IgG secondary antibody (Vector BA-9200, 1:200 [10 μg/ml]) was diluted in 10% guinea pig serum with 1% BSA and applied to sections for 30 min. Antibody signal was amplified using ABC (Vector) for 30 minutes and visualized with DAB with nickel (3,3′-diaminobenzidine tetrahydrochloride, Vector) to create a black precipitate. This process from protein block onward was repeated with anti-pan cytokeratin (2 hour primary at room temperature) and visualized with DAB excluding nickel to form brown precipitate. Nuclei were counterstained using hematoxylin and coverslips mounted using DPX mounting solution (Millipore). Sections were imaged using the Zeiss Axioscan Scanning Microscope and Zen Imaging Software v.3.5 at 40x magnification. N numbers: Control: Dams: n=2, placentas: n=7; MNR: Dams: n=3, placentas: n=7; MNR + IGF1: Dams: n=2, placentas n=8

### Morphometry Analysis

Morphometric analysis was performed using 10 random 40x fields of view of double labeled IHC. Trophoblast and maternal blood space volumes were calculated using point counting with an isotropic L-36 Merz grid via Fiji ImageJ v.2.14.0/1.54f [36]. Counting was performed as described in Roberts et al [22]. Placental capillary number was calculated by counting capillary lumen per field of view, and an average was calculated for each placenta.

### Statistical Analysis

All data was analyzed using SPSS Statistics 29. Distribution assumptions were checked with a Q-Q-Plot. Statistical significance was determined using generalized estimating equations with gamma log link as the distribution/link function. Dams were considered the subject, with diet and nanoparticle-mediated *hIGF1* exposure and direct injection treated as main effects. Gestational age, sex, and litter size were set as covariates. In the sham treated Control and MNR groups, direct placental injection had no effect for any outcomes measured and was therefore removed as a main effect in these groups. Fetal sex-specific differences were not found amongst any of the data and therefore, outcomes for placentas from male and female fetuses were grouped together. Statistical significance was considered at P≤0.05. Bonferroni post hoc analysis was performed for statistical significance. Results are reported as estimated marginal means ± 95% confidence interval.

## RESULTS

Maternal physiological outcomes, fetal growth outcomes, and placental weight/efficiency have been previously published [23]. Details of mating, pregnancies, and fetal weight restoration with repeated *hIGF1* nanoparticle treatment were previously described[23]. One fetus in each pregnancy received a direct nanoparticle-mediated *hIGF1* or sham injection into its placenta, while the remaining fetuses of the litter received indirect exposure from residual circulating nanoparticle circulating in the maternal system. At time of sacrifice, it was determined that all placentas that received repeated direct injection of *hIGF1* nanoparticle treatment were male. As fetal sex cannot be determined at time of first injection and this random determination occurred, data from the MNR + IGF1 (Direct Injection) group only includes males, while all other groups include both sexes. No changes were seen in gross morphology of the placentas between any groups, nor hemorrhages or gross anomalies (Figure 1).

**Figure 1.**
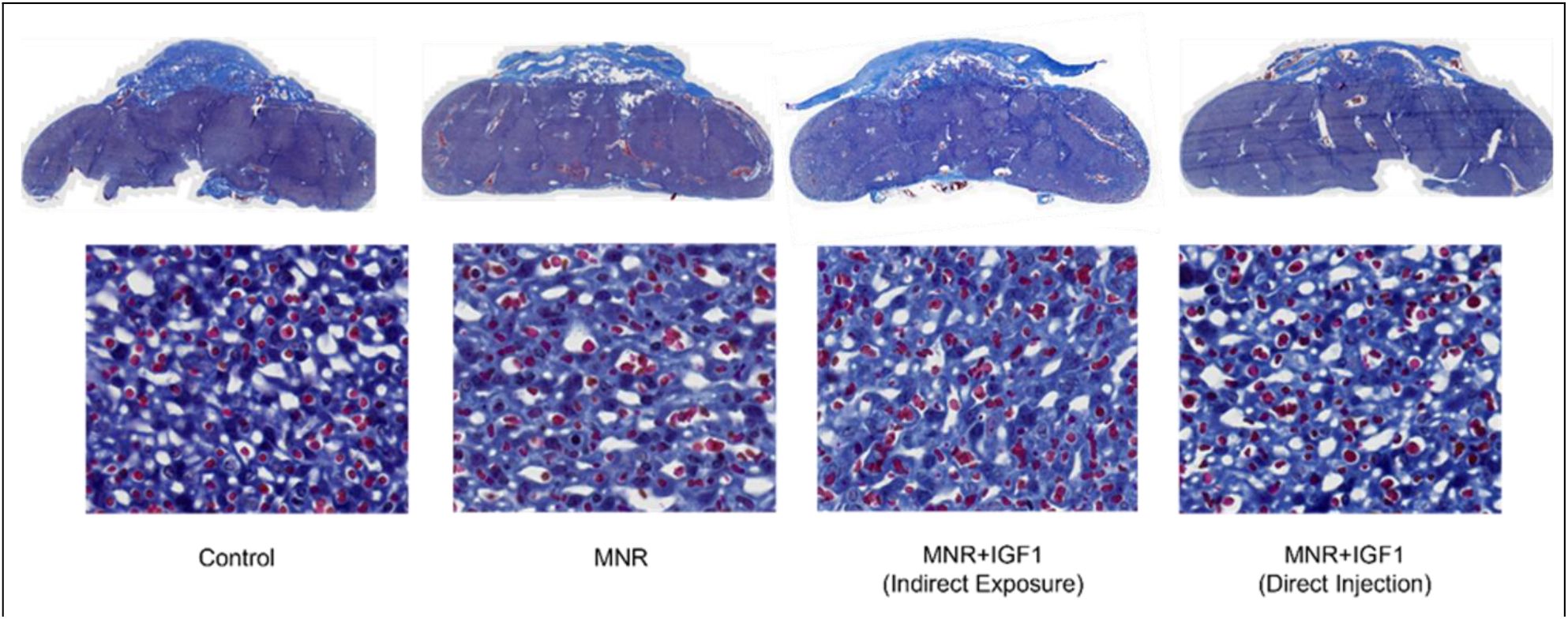
Effects of maternal nutrient restriction (MNR) and repeated *hIGF1* nanoparticle gene therapy (MNR + *hIGF1*) at end of term on gross placenta morphology. Whole scan trichrome images of placentas and sub-placenta/decidua (top), and 40x high resolution images of labyrinth area (bottom).

Using DiceCT methodology and volume graphics machine learning, we 3D rendered Control, MNR, and MNR + IGF1 placentas to determine the structure of their macrovasculature (Figure 2A). While this methodology does not differentiate between macro- and micro-structures, we obtained a resolution limit of >∼13 µm when scanning and therefore could only identify the macrovasculature as seen in the rendering of the solid placenta volume and macrovasculature segmentation (Figure 2B).

**Figure 2.**
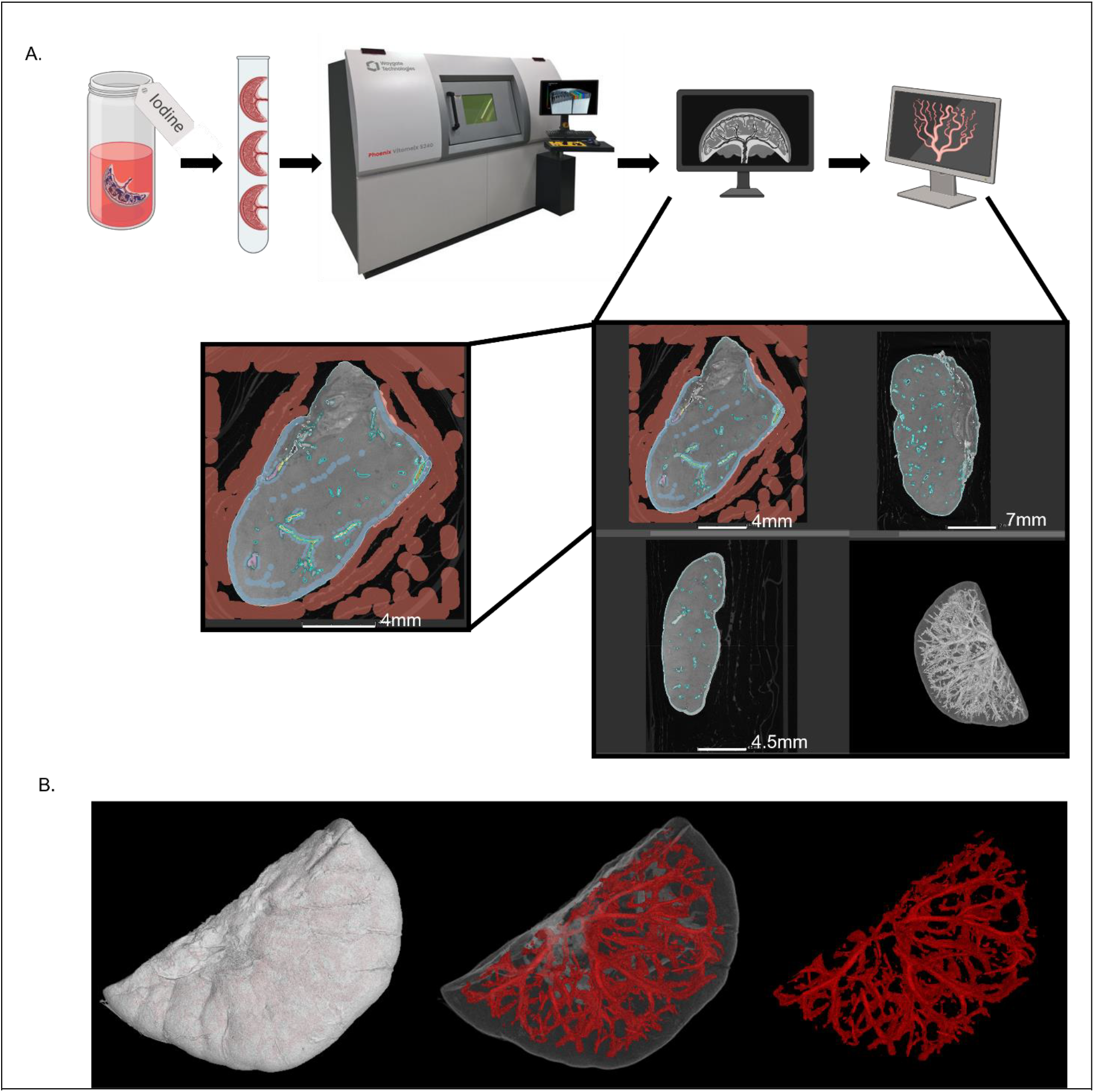
Schematic of DiceCT methodology used to create 3D rendering of placenta macrovasculature structure. **A**. Placentas were stained with iodine, packaged for stability during scanning, scanned with NanoCT to create 2D tiff stack images, and 3D rendered. VG Studio machine learning tool was used for vasculature segmentation for macrostructure modeling. Zoomed image displays machine learning training where red indicates “air/packaging” around the specimen, sky blue indicates “placenta”, purple and yellow indicate “macrovasculature”, and teal identifies areas of learned segments. **B**. Whole volume placenta 3D rendering, Macrovasculature shown within the placenta, and segmented macrovasculature

Vasculature 3D segmentation highlighted the complex placenta macrovasculature of maternal and fetal circulation, showing natural variation within each experimental group (Figure 3). The size of the macrovasculature, both volume (how much space the blood can occupy) and surface area (how large the external surfaces of the vessel is), dictate how much and with what force blood flows through these structures and can provide insights into force on these vessels, back pressure. To elucidate changes in size of the macrovasculature, we quantified the volume and surface area of each segmented macrovasculature. Total volume of the CT segmentation of the macrovasculature decreased in MNR + IGF1 (Indirect Exposure) and MNR + IGF1 (Direct Injection) from MNR (p<0.001, p<0.05), but no group was different from Control (Figure 4A). Surface area of segmented macrovasculature increased in MNR from Control (p<0.05) (Figure 4B). Surface area decreased in MNR + IGF1 (Indirect Exposure) and MNR + IGF1 (Direct Injection) placentas compared to MNR (p<0.01, p<0.05), with no differences from Control.

**Figure 3.**
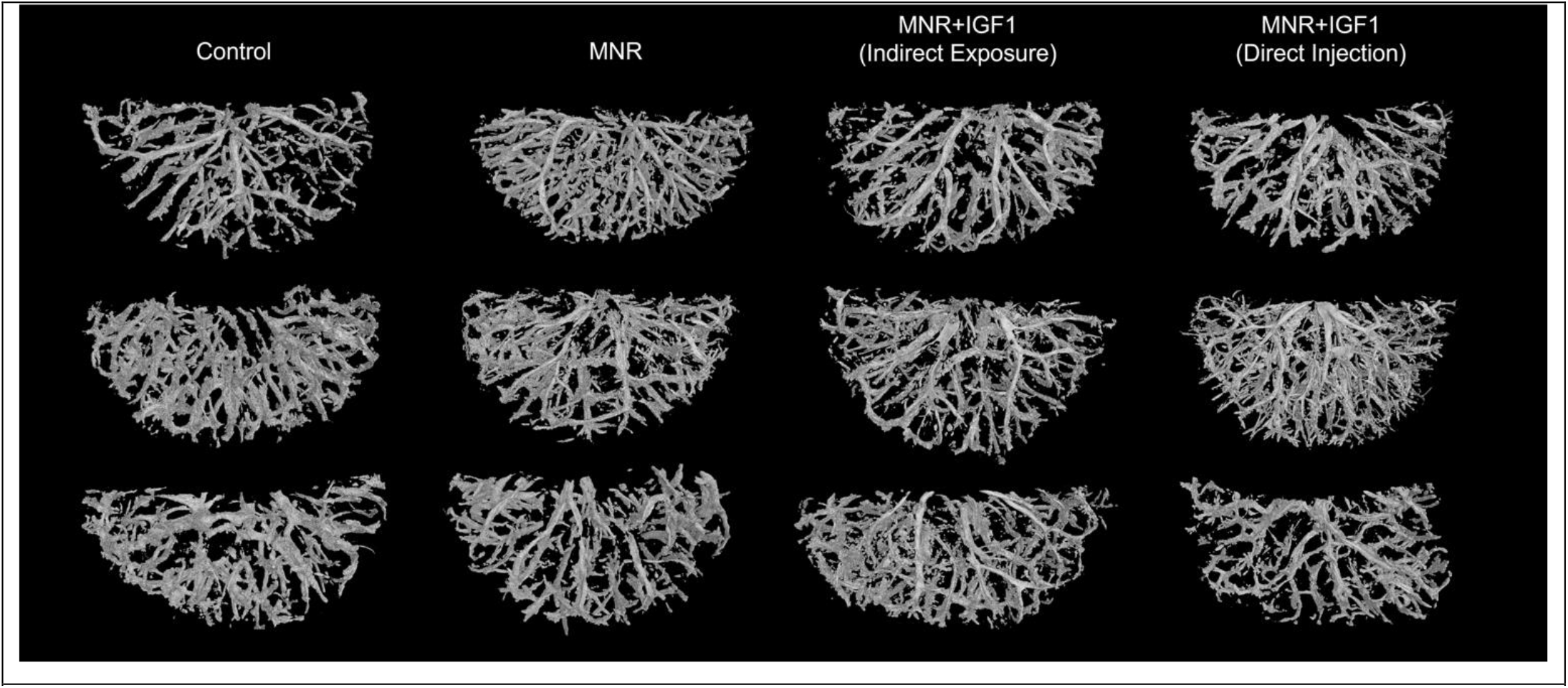
Segmentation of Control, maternal nutrient restriction (MNR), and repeated *hIGF1* nanoparticle gene therapy (MNR + *hIGF1*) placental vasculature. Representative images of segmented placenta macrovasculature of male fetuses.

**Figure 4.**
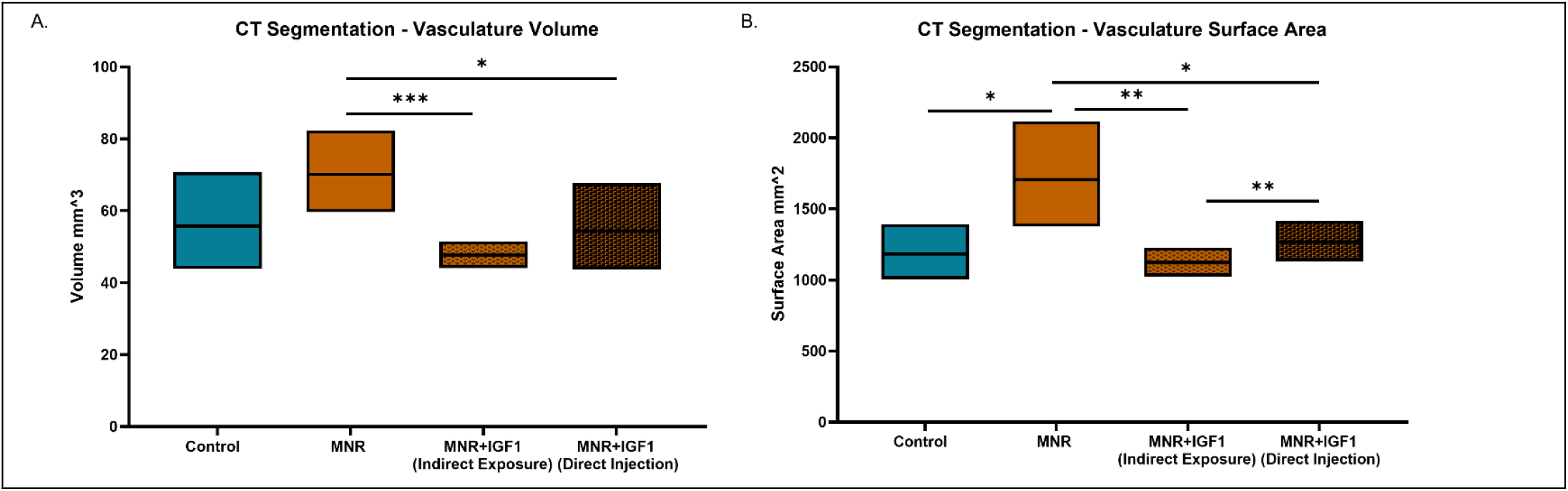
Effects of maternal nutrient restriction (MNR) and repeated *hIGF1* nanoparticle gene therapy (MNR + *hIGF1*) at end of term on placental macrostructure. **A**. Vasculature volume of placenta macrostructure decreased in MNR + IGF1 (Indirect Exposure) and MNR + IGF1 (Direct Injection), but no group was different from Control. **B**. Surface area of placenta macrostructure was increased in MNR compared to Control, MNR + IGF1 (Indirect Exposure), and MNR + IGF1 (Direct Injection). Surface area of MNR + IGF1 (Direct Injection) placentas increased from MNR + IGF1 (Indirect Exposure), but neither differed from Control. Control: Dams: n=2, placentas n=6, MNR: Dams: n=3 female, placentas n=9, MNR + IGF1: Dams: n=3, placentas n=9. Data are estimated marginal means ± 95% confidence interval. *P≤0.05; **P≤0.01. ***P≤0.001

As we could not define maternal from fetal circulation in our macrovasculature CT segmentation, we identified each circulation using IHC to analyze the microvasculature of the labyrinthine exchange area. Immunohistochemistry against Cytokeratin allowed identification of labyrinthine trophoblast and vimentin to identify stroma and placental capillaries. The structure of the labyrinth showed a lack of defined maternal blood spaces in MNR compared to Control (Figure 5). Placental capillaries were also collapsed with indistinct lumens. This implied an increase in interhaemal distance compared to Control whereas MNR + IGF1 (Indirect Exposure) and MNR + IGF1 (Direct Injection) both appeared similar to Control with clear lumens and open maternal blood space.

**Figure 5.**
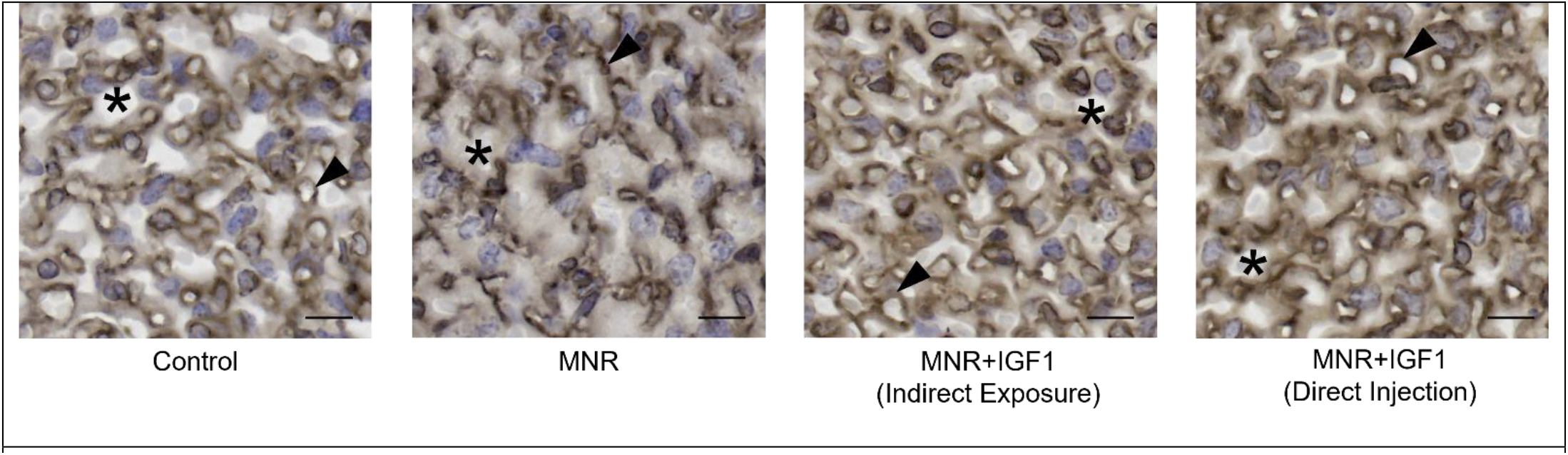
Effects of maternal nutrient restriction (MNR) and repeated *hIGF1* nanoparticle gene therapy (MNR + *hIGF1*) at end of term on placental exchange area. Representative images of immunohistochemistry for cytokeratin and vimentin showing altered placenta exchange area. MNR placentas show a lack of defined structure without clear placental capillary lumens or open maternal blood spaces compared to Control, MNR + IGF1 (Indirect Exposure), and MNR + IGF1 (Direct Injection). Brown: cytokeratin, trophoblast. Black: vimentin, endothelial cells. Stars represent maternal blood spaces; arrow heads represent placental capillaries. Scale bar = 10 μm. Control: Dams: n=2, placentas: n=7; MNR: Dams: n=3, placentas: n=7; MNR + IGF1: Dams: n=2, placentas n=8

Placental capillary count was decreased in MNR compared to Control (Figure 6). MNR + IGF1 (Indirect Exposure) and MNR + IGF1 (Direct Injection) placental capillary count were both increased from MNR, back to Control levels. For greater clarity and identification of large macrovasculature vessels and microvasculature exchange area vessels within the same image/sample, these areas are identified in a representative Control IHC image (Figure 7). Images of the placental disc identify macrovasculature vessels, microvasculature exchange areas, and interobulum areas.

**Figure 6.**
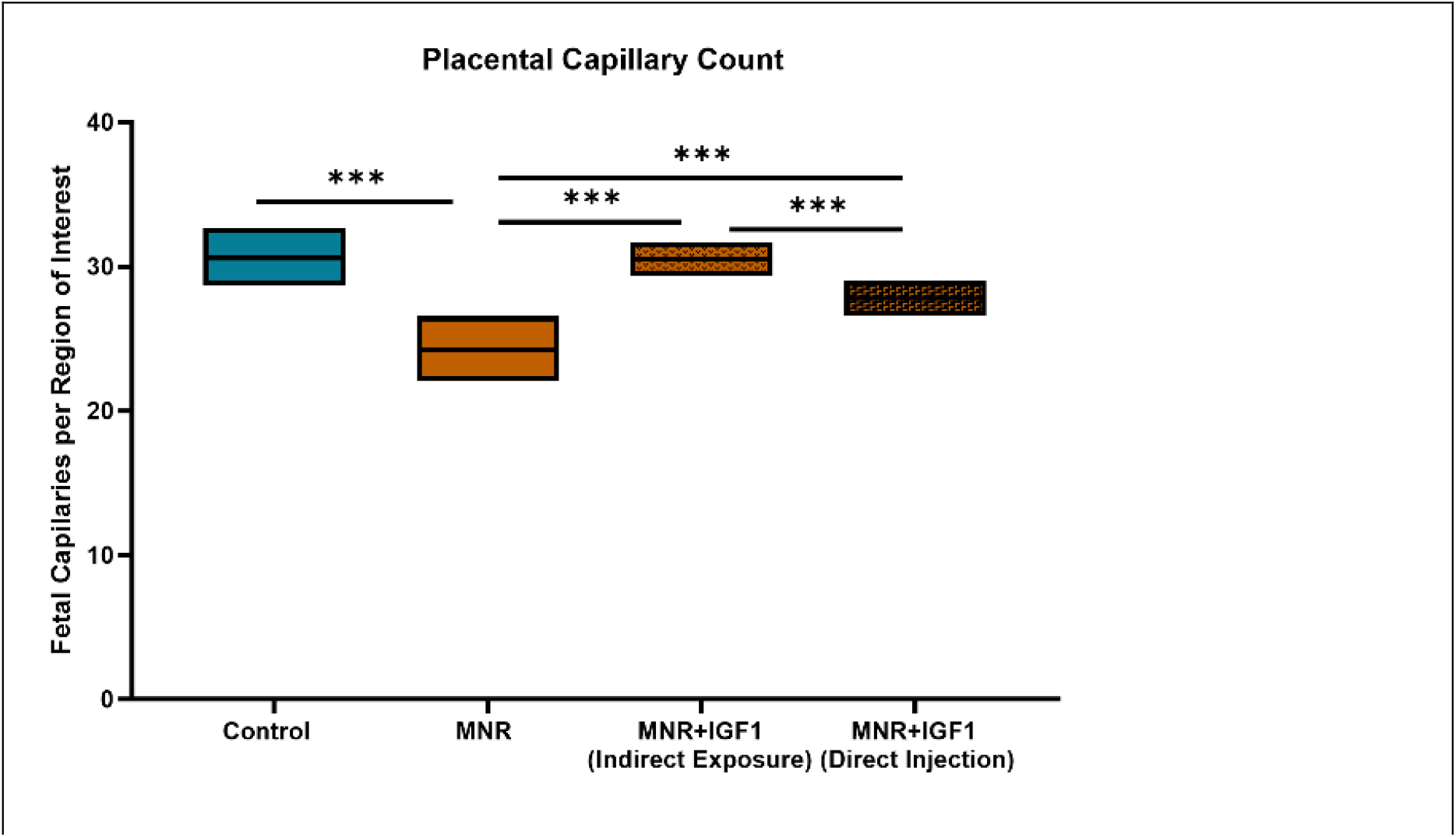
Effects of maternal nutrient restriction (MNR) and repeated *hIGF1* nanoparticle gene therapy (MNR + *hIGF1*) near term on labyrinth capillary count. Placental capillary number decreased in MNR compared to Control MNR + IGF1 (Indirect Exposure), and MNR + IGF1 (Direct Injection). Control: Dams: n=2, placentas: n=7; MNR: Dams: n=3, placentas: n=7; MNR + IGF1: Dams: n=2, placentas n=8. Data are estimated marginal means ± 95% confidence interval. *P≤0.05; **P≤0.01. ***P≤0.001

**Figure 7.**
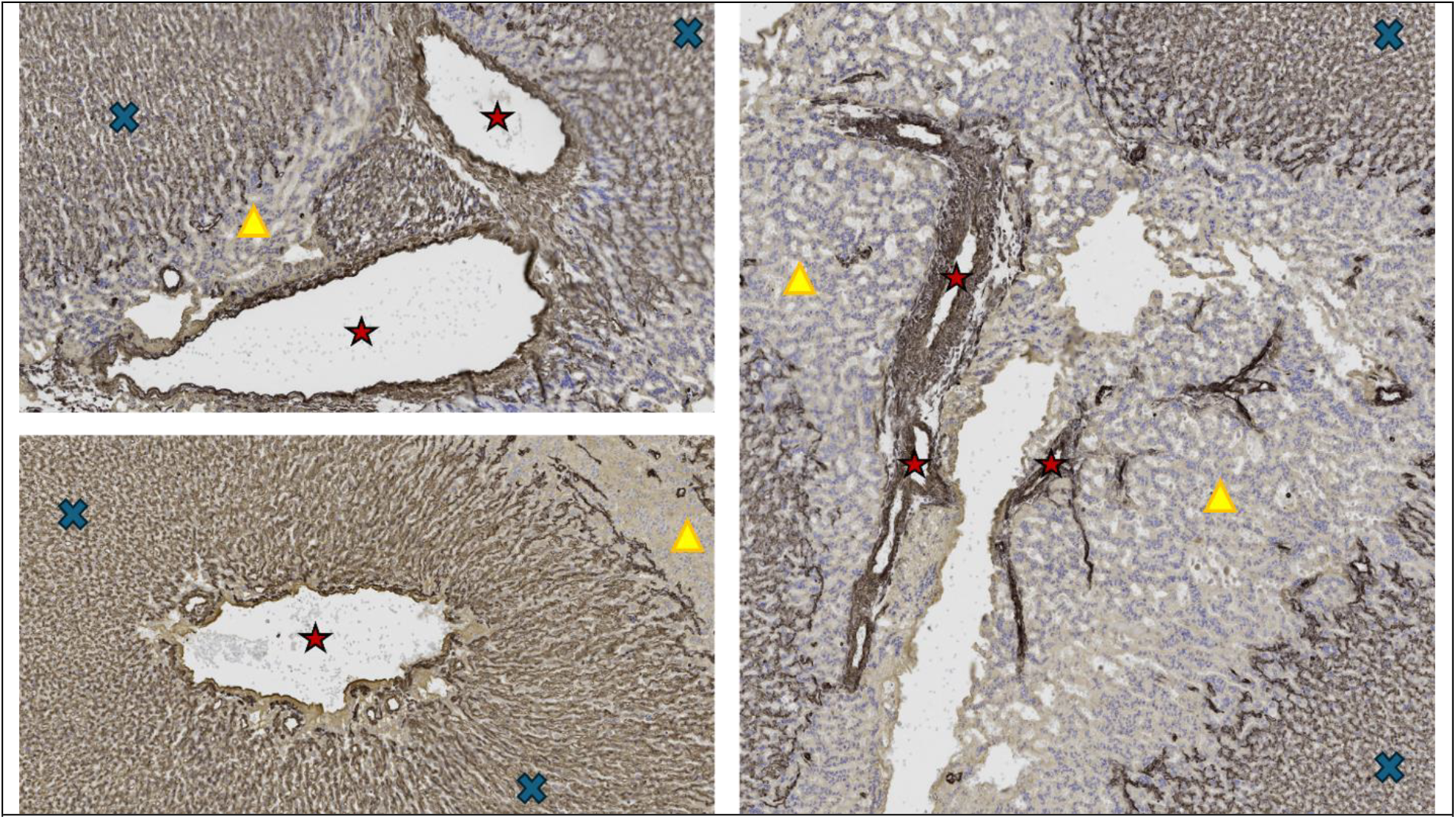
Macrovasculature vessels and microvasculature exchange regions of the placenta. Representative images of immunohistochemistry for cytokeratin and vimentin of Control Placentas. Red stars represent macrovasculature vessels, blue Xs represent microvasculature exchange areas of the labyrinth, yellow triangles represent interlobulum.

## DISCUSSION

There are currently no treatments for fetal growth restriction or placental insufficiency. We have previously shown our nanoparticle-mediated *hIGF1* gene therapy corrects fetal growth, placental weight/efficiency, fetal blood glucose, and maternal and fetal cortisol [23]. In our present study, we aimed to determine the impact of this *hIGF1* treatment on placenta structure to elucidate changes that occur in the placenta labyrinth, leading to better fetal outcomes. Overall, this data demonstrates the aberrant placental structure occurring with MNR at the end of pregnancy is corrected with our nanoparticle-mediated *hIGF1* treatment.

Doppler ultrasound is frequently used in diagnosis of FGR in humans to assess the blood flow and resistance in the placenta [37-39]. If resistance is high in the uterine arteries or blood flow is low in the umbilical artery, the fetus is at risk for FGR [37, 39, 40]. P(z) value is a theoretical representation of back pressure in the uteroplacental circulation, and the pulsatility index (PI) assesses the circulatory status or resistance to flow in the vasculature [40]. From these values we know that both maternal and fetal circulation are disrupted in cases of FGR. While we are unable to get a clear idea of vasculature remodeling across human pregnancy for ethical reasons, animal models offer an opportunity for clearer insights into the structural causes leading to changes in P(z) and PI with placental insufficiency [41].

Using DiceCT, 3D models of the placenta macrovasculature were segmented in our guinea pig model. Quantification of this segmentation revealed increased surface area of the macrovasculature in MNR placentas. From human doppler measures of both umbilical and uterine arteries, we know that back pressure is a common problem in FGR placentas [2, 38, 40, 42, 43]. Increase in surface area of a vessel is generally a response in an effort to lower blood pressure [44]. Forced vasodilation, leading to increased surface area of the vessel, results in decreased blood pressure by distributing the force of the blood over a larger area, reducing the back pressure against the vessel walls, which likely explains the increase in surface area in MNR placentas [43-45]. Volume of the macrovasculature, though not significant, showed the same trend to surface area. As blood volume within a vessel increases, like that in cases of back pressure, the pressure exerted on the vessel walls rises, once again potentially explaining this increase in volume [44]. Repeated nanoparticle-mediated *hIGF1* treatments, however, resulted in surface area and volume no different from Control. We hypothesize increase in vasculature size in MNR may be due to the increased back pressure of the placental circulation as seen in human cases when the P(z) value increases in FGR, while placentas receiving repeated *hIGF1* treatment did not have aberrant back pressure and therefore no vasodilation of the macrovasculature.

While we cannot differentiate the maternal from fetal circulation in our macrovasculature CT segmentation, investigating the microvasculature (∼5-10 µm vessels) of the labyrinth exchange area identified reduced numbers of placental capillaries in MNR significantly decreased from Control placentas. Optimal oxygen diffusion from the maternal to fetal circulation depends on the thickness of tissue, therefore the smaller the interhaemal distance, the easier for oxygen to diffuse [46]. The lack of defined large maternal blood spaces and collapsed placental capillaries separated by larger trophoblast/stroma area in the MNR IHC allude to an increased interhaemal distance compared to Control and MNR+IGF1. At mid-pregnancy there was no change in this interhaemal distance with MNR yet, but *hIGF1* treatment reduced this distance [30].

As gestation progresses, branching of capillaries is essential for placental development to deliver both the increasing amount of nutrients necessary for the exponential growth of the fetus and ensure optimal oxygen transfer to fetal hemoglobin [45, 47, 48]. Decreased number of placental vessel lumens in MNR placentas suggests decreased angiogenesis from Control. With MNR as previously demonstrated by Roberts et al and recapitulated in our current study, placental architecture shows an overall deficit in the exchange area of the placenta elucidating one of the potential main causes of decreased fetal growth/placental insufficiency [22]. With repeated *hIGF1* treatment, however, labyrinthine structure including maternal blood spaces and placental capillaries were restored to Control levels indicating improved angiogenesis and exchange potential.

While previous studies of the guinea pig placenta structure published changes with MNR, we are the first to employ the DiceCT method to reconstruct the guinea pig placenta macrovasculature [22, 49-52]. As previously mentioned, this method is frequently used on museum specimens and/ or whole animal tissues to visualize soft tissue and anatomic organization of organ systems [32, 34]. Limitations of this novel approach include staining times and concentrations dependent on type of tissue (easily corrected with water baths or higher concentration of Lugol’s iodine) and machinery/software improvements. While we were able to successfully segment the macrovasculature of these placenta, a higher resolution scan on the current GE CT, or an alternative CT such as the Ziess Versa 620 XRM, gives resolution down to 50 nanometers and a more precise segmentation that include the microvasculature exchange area and the macrovasculature may be possible in the future. This tool is likely to improve with each software update and version release, and we recommend it for complex structures such as the placenta vasculature. Interestingly, in contrast to other measurements we published, the structural differences and remediation reported here did not vary based on fetal sex. In future studies, our goal is to include doppler measures for uterine and umbilical artery flow to elucidate the similarities in blood flow within the guinea pig placenta to human cases of FGR and assess the effects our *hIGF1* treatment.

In the present study, we show changes in the placental macrovasculature and microvasculature exchange area of the labyrinth in MNR placentas that recapitulate human placental insufficiency. We also show the ability of our novel nanoparticle-mediated *hIGF1* gene therapy to improve placental architecture for proper fetal growth.

## Notes

### Competing Interest Statement

The authors have declared no competing interest.

